# TRB3 augments IL1β-TLR4 signaling by engaging Flightless-homolog 1

**DOI:** 10.1101/2022.10.09.511391

**Authors:** Sumati Gonuguntla, Rohan K. Humphrey, Akshita Gorantla, Ergeng Hao, Ulupi S. Jhala

**Affiliations:** Pediatric Diabetes Research Center, University of California San Diego, La Jolla CA 92037

**Keywords:** Tribbles, FLII, MLK3, IKK, NFkB, Interleukin1, MyD88, Toll-Like Receptor, pancreatic β cells, inflammation, TRIB3, pseudokinase, Tribbles

## Abstract

Signaling via IL1β and TLR4 receptors (IL1R-TLR4) plays a crucial role in cytokine and fatty acid-induced beta cell inflammation, in type 1 and type 2 diabetes respectively. IL1R-TLR4 share signaling mechanisms via a common, cytoplasmic, toll-like-receptor domain to activate proinflammatory JNK and IKK kinases. We have previously reported that in response to IL1β, pancreatic islets isolated from TRB3 knockout (TRB3KO) mice show attenuated kinetics of activation for MAP3K MLK3, and JNK stress kinases. Here we report that similar to MLK3 and JNK, TRB3KO islets also show a decrease in amplitude and duration of IL1β/LPS-stimulated TAK1 and IKK phosphorylation. Thus, loss of TRB3 attenuates both pathways critically required for a full-blown, cytokine-inducible, proapoptotic response in beta cells. TRB3KO islets display a sharp decrease in cytokine-induced beta cell death, accompanied by a decrease in select downstream NFkB targets, most notably, inducible Nitric Oxide Synthase (iNOS/NOS2), a well-characterized mediator of beta cell dysfunction and death. In order to better understand the molecular basis of TRB3-enhanced IL1R-TLR4 signaling, we interrogated the TRB3 interactome and identified Flightless-homolog 1 (Fli1), an immunomodulatory, actin-binding, leucine-rich-repeat protein, as a novel TRB3-interaction factor. TRB3 binds and disrupts Fli1-dependent sequestration of MyD88, thereby increasing availability of this proximal adaptor to participate in IL1R-TLR4 signaling. Fli1 forms a multiprotein complex that can disconnect IL1R-TLR4 from MyD88, resulting in a brake on assembly of downstream signaling complexes. By interacting with Fli1, TRB3 lifts the brake on IL1R-TLR4 signaling to augment the proinflammatory response in beta cells.

## Introduction

Beta cell inflammation contributes to the etiology of both type 1 and type 2 diabetes (T1D and T2D). However, the rate and degree of beta cell inflammation in T1D versus T2D are distinctly different. The strength and nature of signal-dependent physiologic responses are determined by spatial and temporal dynamics of signal activation (1). In turn, the dynamics of signaling are strongly influenced by the scaffolds and adaptors that molecularly wire the signaling machinery (2), highlighting the need for a more nuanced understanding of pro-inflammatory signaling in the beta cell. TRB3 is a stress-induced pseudokinase whose function is unmistakably associated with a number of pathologies, including obesity, diabetes and Parkinson’s disease (3,4). At the cellular level, TRB3 has been characterized as an inducer of ER stress, insulin resistance and apoptosis (3,5,6). However, the molecular mechanism/s by which TRB3 exerts its effects are less well understood. The gap in information is partly because TRB3 is a pseudokinase that lacks critical motifs required for ATP binding and transfer (7), resulting in absence of a clear readout of TRB3 molecular action. We have previously reported that in beta cells, IL1β rapidly induces phosphorylation if MAP3K MLK3 to in turn, stimulate JNK stress kinase. pMLK3 also binds and stabilizes TRB3 protein leading to a feed forward mutually stabilizing effect on components of the JNK module including JIP1, MLK3 and TRB3(8). Inhibition of MLK3 protects beta cells from caspase-dependent cell death in the early phases of inflammation, but has no effect on the activation of IKK-NFkB pathway (5), the second proinflammatory signaling arm required for a full and robust inflammatory program. Here we provide evidence to show that in addition to MLK3-JNK signaling, TRB3 also regulates beta cell inflammation by attenuating TAK1-IKK phosphorylation and subsequent NFkB-mediated expression of key genes that drive beta cell inflammation. Although TRB3 has been reported to function as an allosteric regulator of multiple kinases (3,9,10), we were unable to identify clear and direct, binary interactions between TRB3 and kinases that regulate IKK-NFkB pathway in the beta cell. Here we describe a novel interaction between TRB3 and the LRR (Leucine Rich Repeat) protein Flightless Homolog-1(Fli1). Fli1 is an actin-binding molecular scaffold protein (11,12), which can cooperatively enhance, or competitively inhibit cellular inflammation (13). TRB3 engages Fli1 in response to cytokine stimulation, to augment IL1β and LPS signaling in the beta cell. Our results suggest that TRB3 can function as a scaffold protein and fine-tune kinase activation for stimulus-dependent activation of the inflammatory response.

## Results

### TRB3 is required for optimal IL1β/TLR4 mediated signaling

The pseudokinase TRB3 is induced by multiple stress stimuli including cytokines, and agents that induce ER stress (5,6). In the beta cell, MAP3K-JNK, and IKK-NFkB signaling pathways are the most potent mediators of cytokine-dependent inflammation and decline in beta cell survival (14,15). We have previously reported that in beta cells, the pseudokinase TRB3 modulates the strength of JNK activation by stabilizing a cytokine-activated signaling module consisting of MLK3, MKK7, JIP1, and JNK1/2 (5,10). As previously reported (10), and shown here in Figure 1A, such a role for TRB3 is associated with optimal activation of MLK3 and sustained activation of JNK in islets from WT mice, and both are attenuated in pancreatic islets from TRB3KO mice. These results prompted us to examine whether TRB3 could similarly modulate activation of the IKK-NFkB pathway. IL1β stimulation of the canonical IKK-NFkB signaling pathway requires the activation of TAK1 kinase to activate downstream gene expression (16,17). MLK3 and TAK1 are both activated by signal-dependent ubiquitination and autophosphorylation, downstream of the IL1β receptor (16,18,19). We therefore examined IL1β-mediated phosphorylation of TAK1 and IKK in islets from WT and TRB3KO mice. Isolated islets from WT TRB3KO mice were stimulated with ILβ in a time dependent manner, and islet extracts used for Western blotting. As seen in Figure 1B and 1C, when compared to WT islets, TRB3KO islets showed 50-60% lower induction of both TAK1 and its downstream target IKK. In addition to direct activation by cytokines, beta cell inflammation can also be activated by lipotoxicity, principally induced by saturated fatty acids and ceramides (reviewed in (20,21). Studies show that although saturated fatty acids do not directly engage TLR4, the lipotoxic inflammatory response requires TLR4 activation (22,23). TLR4 signaling pathway overlaps significantly with that of IL-1β, including recruitment of downstream adaptor MyD88 and effector TRAF6 (24,25). We therefore examined whether islets from WT and TRB3KO mice exhibited a difference in their response to TLR4 stimulation. As seen in Figure 1D stimulation of islets with TLR4-specific lipopolysaccharide (LPS) also showed marked attenuation of the TAK1-IKK signaling module in TRB3KO islets. Interestingly, compared to IL1β, stimulation of TLR4 showed delayed activation of TAK1 and IKK. In islets, JNK was not activated in response to LPS stimulation (not shown).

**Figure 1:**
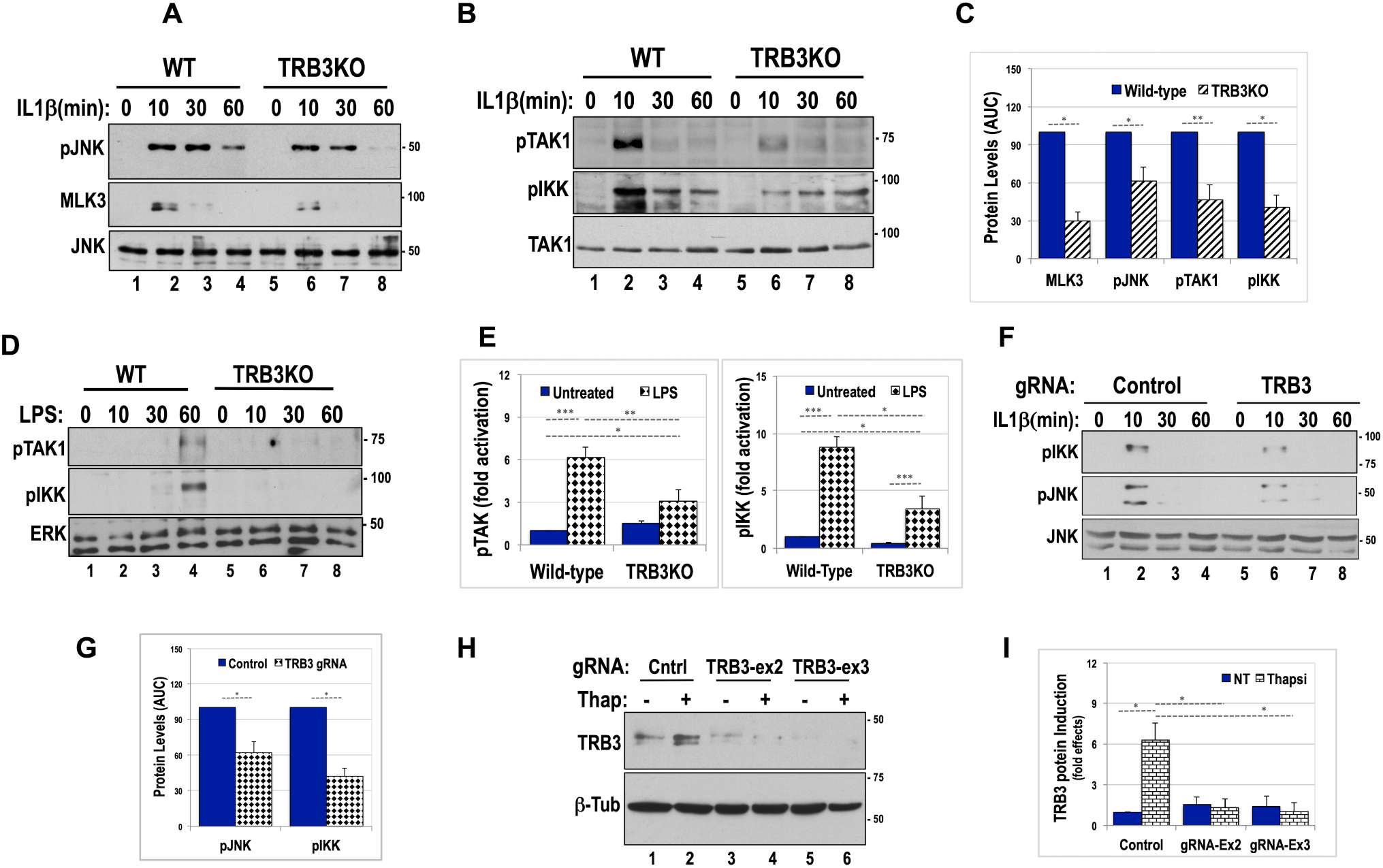
Loss of TRB3 leads to attenuated activation of JNK and IKK in islets and Min6 cells: **A)** Wildtype (lanes 1 to 4) and TRB3KO (lanes 5 to 8) islets were treated with 20ng/ml of IL-1β for the indicated times and lysates were subjected to western blotting using anti-MLK3 and phospho-JNK antibodies. Total JNK protein levels are shown. **B)** Wildtype (lanes 1 to 4) and TRB3KO (lanes 5 to 8) islets were treated with 20ng/ml of IL-1β for the indicated times and lysates were subjected to western blotting using anti-phosphoTAK1 and phospho-IKK antibodies. Total TAK1 protein levels are shown. **C)** Densitometric analyses of time-dependent JNK, IKK, and upstream MAP3Ks MLK3 and TAK1 were graphed and total kinase induction/activity over time assessed by calculating area under the curve (AUC). Average values from three independent time course experiments are presented as percent fold effects of control-WT samples. Two-way ANOVA was performed to compare WT and TRB3KO (*\* p<0.05*). **D)** Wildtype (lanes 1 to 4) and TRB3KO (lanes 5 to 8) islets were treated with 5ng/ml of TLR4 receptor-specific LPS for the indicated times, and lysates were subjected to western blotting using anti-phosphoTAK1 and phospho-IKK antibodies. Total ERK protein levels are shown as control. **E)** Graph shows fold-peak activation of LPS-stimulated kinase (at 60 mins) normalized to unstimulated WT control signal density (*\* p<0.05, **<0.005, *** <0.0001*). Data shown represents densitometric analysis of three independent experiments, each using islets pooled from multiple mice. **F)** Min6 cells electroporated with control (lanes 1-4) or TRB3 gRNA (lanes 5-8), were treated with 20ng/ml IL-1β for the indicated times and lysates subjected to western blotting using anti-phospho-IKK and phospho-JNK antibodies. Total JNK protein levels are shown. Graph shows average, IL-1β-induced kinase activation from TRB3 gRNA expressing cells. Data presented is calculated expressed as percent of control within each experiment (\**p<0.05)*. **H)** Efficacy of TRB3 knockdown was tested using Min6 cells electroporated with control (lanes 1-2) or gRNA constructs targeted to exon 2 (lanes 3-4) or exon 3 (lanes 5-6) of the mouse TRB3 gene. TRB3 expression was induced using thapsigargin (400 nM for 5 hours) as shown. **I)** Quantification of TRB3 protein levels from Figure 1H represent an average of three independent experiments *(*p<0.05)*. Unless specified, data presented in bar graphs show average fold effects from three independent experiments, calculated by normalizing all values to WT control. In each case, mean values +/− SD of three experiments are presented.

Next, we examined whether short-term loss of TRB3 could similarly impact activation of IKK and JNK in Min6 insulinoma cells. For knockdown of TRB3, we used the CRISPR-Cas9-Nickase system, to target two independent guide RNAs, directed to two different exons of nascent TRB3 mRNA. gRNA design to suppress TRB3 induction is shown in Supporting Information figure S1. Nucleofection of Min6 cells showed the efficacy with which each gRNA suppressed stress-induced expression of TRB3 protein in Min6 cells (Figure 1H). Using TRB3 targeted gRNA, we observed an attenuation of IL-1β dependent JNK and IKK activation in Min6 cells. These results closely parallel the results obtained from TRB3KO islets. Total JNK was used as loading control (Figure 1F).

### Loss of TRB3 attenuates NFkB-dependent gene activation

Studies have demonstrated that activation of IKK and it’s downstream effector NFkB are crucial for cytokine-induced inflammatory damage of beta cells (26–28). We examined whether TRB3 expression is required for optimal NFkB-dependent gene activation. Using a NFkB response element-driven luciferase reporter in transient transfections, we examined whether co-expression of TRB3 gRNA impacted activity in IL1β-stimulated Min6 cells. As seen in Figure 2A, IL-1β treatment resulted in a 10-12 fold induction of NFkB-luc activation in the presence of gRNA control, while presence of TRB3 gRNA attenuated ILβ-dependent NFkB-luc activation, dropping by roughly 60% compared to control. These data indicated that the attenuation of IKK seen in TRB3-deficient islets and Min6 cells, extends to downstream, NFkB-dependent gene expression. However, a reporter construct driven by multimerized NFkB response elements does not reflect the complexity of promoters that drive endogenous genes, where additional transcriptional activators could reduce/enhance the impact of IKK activation on gene expression. We examined whether loss of TRB3 impacted mRNA expression of known NFkB targets previously implicated in the inflammatory process. Isolated islets pooled from multiple WT or TRB3KO mice, were treated with IL1β as indicated. GAPDH mRNA was used as an unchanging control. Interestingly, loss of TRB3 did not uniformly impact expression of NFkB target genes. Compared to WT islets, IL1β treatment of TRB3KO revealed a small but significant decrease (30%) in IL1β-stimulated expression of IL6 mRNA in TRB3KO islets (Figure 2B). Similarly upon IL1β treatment, mRNA for iNOS (inducible nitric oxide synthase-NOS2) gene was reduced in TRB3KO islets by 30-40% (Figure 2D). By contrast, compared to WT islets, macrophage chemoattractant protein MCP1(CCL2) was strongly downregulated (80%) in IL1β-treatment of TRB3KO islets as described (Figure 2C). No impact was seen on mRNA of IL1β, and TNF-α in these islets (not shown) at any time point. WT islets showed high basal expression of Cycloxygenase-2 (COX2) mRNA, which trended lower approx 20% in TRB3KO islets (not shown). Overall, our results suggest that partial/attenuated IKK activity seen in TRB3KO appears to have a differential effect on downstream NFkB gene expression.

**Figure 2:**
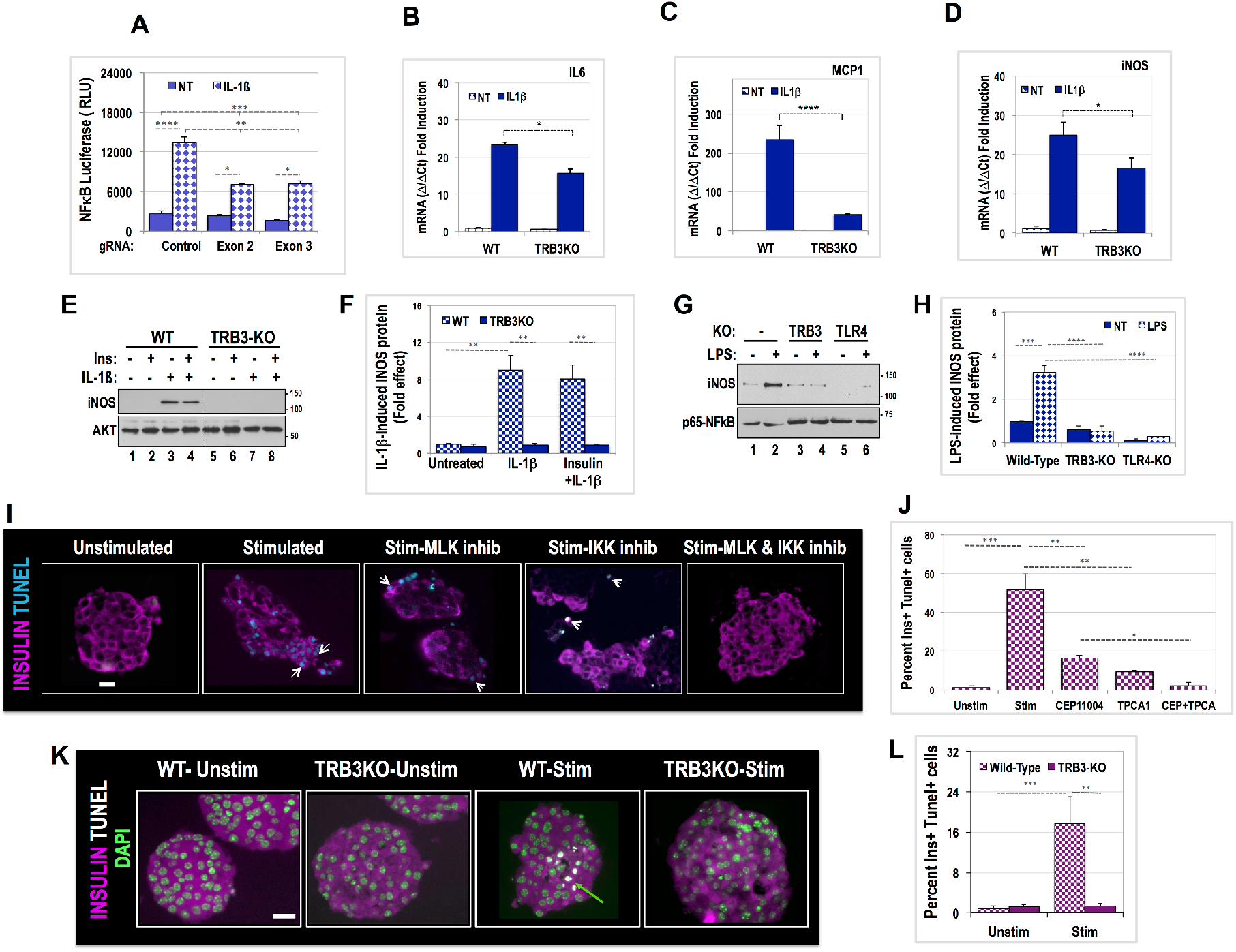
TRB3 is required for optimal activation of NFkB target genes. **A)** Luciferase reporter assay from Min6 insulinoma cells transiently transfected with firefly NFkB-Luciferase reporter in the presence of control gRNA or TRB3 gRNAs. Cells were treated with 20 ng/ml IL-1β (solid bars) or left untreated (hatched bars). Data show results from one of three representative transient transfection experiments. Two-way ANOVA was performed for TRB3 gRNAs against control (*\*\*\*\* p<0.0001, *** p<0.0005, ** p<0.005, * p<0.05*). **B, C, D)** Relative mRNA expression from IL1β-treated primary islets for IL6 (3 hours), MCP1/CCL2 (6 hours), and iNOS (3 hours) gene expression respectively, using RNA isolated from WT and TRB3KO mice. Two-way ANOVA was performed to compare WT and TRB3KO (*iNOS and IL6-*p<0.05, MCP1\*\**\**\*p<0.0001)*. Data represent an average of fold mRNA induction from three independent experiments each performed using mRNA from pooled islets of control and TRB3KO mice. **E)** Western blot examining iNOS expression in protein extracts derived from islets of WT (lanes 1-4) and TRB3KO (lanes 5-8) mice, treated with 20 ng/ml IL-1β and/or 100 nM insulin for 8 hours as indicated. Insulin stimulation preceded IL-1β treatment by 10 minutes. **G)** Western blot for iNOS expression using protein extracts from pooled islets treated with 5 ng/ml LPS for 8 hours. Groups included are WT (lanes 1-2), TRB3KO (lanes 3-4) and TLR4KO mice. **F&H**) Quantification of iNOS protein for 2E and 2G was performed using densitometric analysis of three independent experiments, and represents average protein induction expressed as a fold effect of WT control islets. (Fig 2F *\*\* p<0.005, * p<0.05*) (Fig 2H \*\**\*\* p<0.0001, *** p<0.001*). **I)** Human donor islets were co-cultured with unstimulated or stimulated/activated human splenocytes as described (18). Prior to co-culture, islets were pretreated for 1 hour with MLK inhibitor CEP11004 (500nM), or IKK inhibitor TPCA1 (5μM) as indicated. The islets were harvested and fixed after 30 hours, sectioned and processed for TUNEL staining and immunofluorescence using anti-insulin antibodies. For clarity insulin staining is pseudocolored magenta, and TUNEL staining pseudocolored cyan. White arrow shows apoptotic cells. **J)** Quantification of Insulin expressing cells positive for TUNEL staining, expressed as a % of total insulin+ cells. TUNEL staining was performed simultaneously using sections from four independent co-culture experiments. Data were analyzed using one-way ANOVA (*\*\*\* p<0.0005, ** p<0.005 * p<0.05*) **K)** Islets from WT and TRB3KO mice were co-cultured with immune-activated splenocytes from WT mice for 24 hrs and processed as described in Figure 2I. Insulin stain was pseudocolored magenta, TUNEL staining is pseudocolored white and indicated by an arrow. Nuclei were counterstained with DAPI. **L)** Quantification of insulin/TUNEL double+ cells expressed as a % of total, insulin-expressing cells from multiple islets using sections from three independent experiments. (*\*\*<0.005, *** <0.0001*). Note: Datasets used for statistical analysis in Fig. 2L have been previously published (9).

### Islets of TRB3KO mice fail to show IL1β-LPS induced expression of iNOS protein

Although mRNA levels are an important indicator for understanding role of genes that drive signal dependent cellular behavior, biological effects are determined by protein expression. iNOS protein has been shown to play a pivotal role in cytokine-induced beta cell death (29), prompting us to examine whether reduced iNOS gene expression in TRB3KO islets resulted in decreased protein expression. We found that post-cytokine treatment, iNOS protein is maximally induced by 8 hours, persisting upto 12-15 hours (not shown). We therefore treated pooled islets from each group with IL1β for 8 hours and as shown in Figure 2E, Western blotting of islet extracts showed a robust induction of iNOS in WT islets, which was almost entirely abolished in TRB3KO islets. The proapoptotic role of TRB3 in many cell types has been attributed to TRB3-dependent inhibition of AKT prosurvival kinase (3). Studies in primary hepatocytes have revealed that AKT activity lowers iNOS expression either directly or indirectly via inhibition of GSK3 (30,31). We therefore examined whether the steep decrease in iNOS protein expression in TRBKO islets could be overcome by pretreatment of insulin. We found that WT islets pretreated with insulin showed a marginal but statistically insignificant decrease in iNOS expression (Figure 2E and F). These data clearly show that the pro-inflammatory function of TRB3 extends well beyond any ability to inhibit AKT in beta cells. Having shown that LPS-induced IKK expression is attenuated in TRB3KO islets, we examined whether iNOS expression was also attenuated by LPS in TRB3KO islets. As seen in Figure 2G, and similar to data for IL-1β, treatment with TLR4-specific LPS upregulated iNOS expression in WT islets but showed no induction in TRB3KO islets. Furthermore, the near total inability of LPS to induce iNOS protein expression in TRB3KO islets matched that seen in TLR4KO islets. Taken together our data show that TRB3 is required for optimal IL1R/TLR4 pathway-induced expression of iNOS, a crucial contributor to cytokine-activated oxidative stress and acceleration of beta cell death (32).

### Pharmacologic inhibition of MLK3 and IKK suppresses cytokine-induced beta cell apoptosis

TRB3KO islets show attenuated expression of two critical pathways in cytokine-induced islet inflammation. We have previously shown MLK3-JNK pathway is activated in the early stages of cytokine-induced beta cell death and pharmacologic inhibition of MLK family of kinases (500nM CEP11004) delays onset of beta cell death by several hours. MLK3-JNK pathway was associated with mitochondrial outer membrane permeabilization (MOMP) via mitochondrial translocation of BAX, a proapoptotic BCL2-family protein (33). Notably, MLK3 inhibition had no effect on IKK-NFkB-dependent target activation in islets. Given that TRB3KO islets display attenuation of both MLK3-JNK and TAK1-IKK pathways, and that IKK-NFkB targets are instrumental in inducing islet dysfunction, we examined whether pharmacologic inhibition of IKK-NFkB activation could supplement the effect of MLK3 inhibition. We also examined whether inhibition of both pathways could inhibit rather than just delay progression of beta cell apoptosis. Although pharmacologic inhibition of kinases is often fraught with non-specific inhibition of additional pathways. The main goal of these experiments was limited, and meant solely to demonstrate proof of principle that both MLK3-JNK and TAK1-IKK pathways contribute to apoptosis using primary islets.

Towards this goal, we improvised and developed an islet culture method that could induce beta cell death at levels observed in type 1 diabetes (5) (see schematic in Supporting Information-Figure S2). We used human donor islets and cadaveric human spleens in this assay as described (18). Primary donor human islets were suspended in transwells and cocultured with plated activated human splenocytes for approximately 30 hours. Splenocytes were activated to elicit a robust immune reaction (see Supporting Information, Table 1) using anti-CD3 and anti-CD28 antibodies, while unstimulated splenocytes served as a control. As shown in Figure 2I, we examined whether inhibition of MLK3 using CEP11004 along with IKK inhibitor TPCA1 was sufficient to rescue beta cells from cytokine activated cell death. A thirty-hour incubation of islets with stimulated splenocytes resulted in widespread cell death (approx. 40%) as judged by strong TUNEL staining which detects DNA strand-breaks associated with apoptosis. Islets incubated with stimulated medium also displayed extensive loss of islet morphology and noticeable loss of insulin from beta cells across islets. Pretreatment of islets with MLK3 inhibitor CEP11004, or TPCA1, a powerful IKK inhibitor were each partially effective at inhibiting widespread cell death (reduced from roughly 40-50% without drugs to approximately 8-15% (Figure 2J). When CEP11004 and TPCA1 were both used, we observed near complete inhibition of cell death, a better retention of insulin, and preservation of islet morphology. It may be noted that BAY-117085, an alternate inhibitor of the TAK1-IKK pathway had a comparable effect on cell death (not shown). Although we cannot completely rule out the off-target effects of each drug, our data show that inhibition of both MLK3-JNK and TAK1-IKK pathways could be sufficient to preserve human beta cell integrity in response to inflammation.

### Genetic ablation of TRB3 confers resistance to cytokine induced beta cell death

Pharmacologic inhibitors are strong drugs that result in near complete inhibition of their targets. Such an inhibition would be well beyond the attenuation of the MLK3-JNK and TAK1-IKK pathways seen in TRB3KO islets. Additionally, drugs have off target effects that could affect the interpretation of results. In an experiment identical to that reported in our previous study (9), we examined whether loss of TRB3 could exert a protective effect on cytokine induced cell death. Again, isolated islets from both TRB3KO and WT mice were incubated with stimulated splenocytes from WT mice, in an assay similar to that used for human islets in Figure 2I. Compared to WT islets, upon co-culture with stimulated splenocytes, TRB3KO islets were largely negative for TUNEL staining (Figure 2K). Interestingly, mouse islets did not reveal the widespread loss of insulin or a breakdown in islet morphology. It is unclear whether a better-preserved islet morphology in mouse stimulated islets is species-specific or due to a far reduced time span between isolation and assay. Nevertheless, taken together our data show that TRB3 expression is required to augment proinflammatory signaling to thresholds high enough for triggering cytokine-stimulated cell death.

### Cytokines enhance TRB3 interaction with the scaffold protein Flightless homolog I (Fli1)

TRB3 structurally resembles kinases but lacks critical residues for catalytic phospho-transfer. Therefore, it mediates its actions by allosterically modulating the function of interacting proteins. The apoptotic potential of TRB3 has been attributed to its ability to interact with and inhibit AKT, a potent prosurvival and pro-growth kinase (3). We have shown that insulin pretreatment (a stand in for AKT-activation) had marginal effects on the expression of damage-inducing iNOS protein, suggesting that TRB3 proinflammatory action likely involves additional cellular mechanisms. In order to get a better understanding of the mechanism by which TRB3 augments IL-1/TLR4-induced IKK-iNOS activation, we examined whether TRB3 interacts with proteins that could augment cytokine/IKK signaling.

Towards this end, we interrogated the TRB3 interactome by expressing Flag-TRB3 in Min6 mouse insulinoma cells followed by standard co-immunoprecipitation assays, proteolytic digestion of immunoprecipitates followed by by mass-spectrometry (see methods). The experiment was repeated two more times using identical conditions. In none of the experiments were we unable to identify any kinase or other protein that could directly boost IKK activation/signaling (not shown). We did however identify two LRR (Leucine Rich Repeat)-family proteins shown to previously regulate LPS signaling in immune cells. The two proteins, Fli1 (Flightless homolog 1) and LRRFIP2 (Leucine Rich Repeat Flightless interacting protein) are scaffold proteins that function by modulating availability of MyD88, the most proximal adaptor in IL1/TLR4 signaling cascade (34,35). Shown in Figure 3A is the protein sequence of murine Fli1, with recovered peptides highlighted in color. In TRB3 immunoprecipitates, we recovered 42 unique peptides mapping to Fli1 protein, that displayed a 2-fold (log-scale) or higher recovery in the presence of TRB3 versus control. The peptides together covered 47% of the 1261 amino acids of Fli1. The Fli1 protein consists of multiple repeats of two independent domains, including the aforementioned LRR in the N-terminus, and the actin-binding gelsolin domain in the C-terminus (11). Leucine-rich repeat (LRR) motifs consisting of 20-30 amino acids are present in a wide variety of proteins; their distinct shape making them versatile motifs for protein interactions (36). We examined whether the LRR domain of Fli1 (HA-tagged Fli1 N-terminal 400 amino acids) could immunoprecipitate (IP) Flag-TRB3 expressed in Min6 cells. Using a co-IP assay followed by western blotting, we found that HA-Fli1-LRR could indeed pull down co-expressed Flag-TRB3, and this binding was strongly induced in the presence of LPS, even as total levels of HA-Fli1-LRR were decreased upon LPS treatment (Figure 3B). We also examined if the LRR domain of HA-LRRFIP2 could immunoprecipitate Flag-TRB3, but we saw no interaction between the proteins (Figure 3B). These results suggest that TRB3 interaction with Fli1 is most likely specific and not a result of generic interaction with proteins bearing LRR motifs. These data also suggest that interaction with LRRFIP2 observed in the interactome is likely to be indirect and potentially via Fli1.

**Figure 3:**
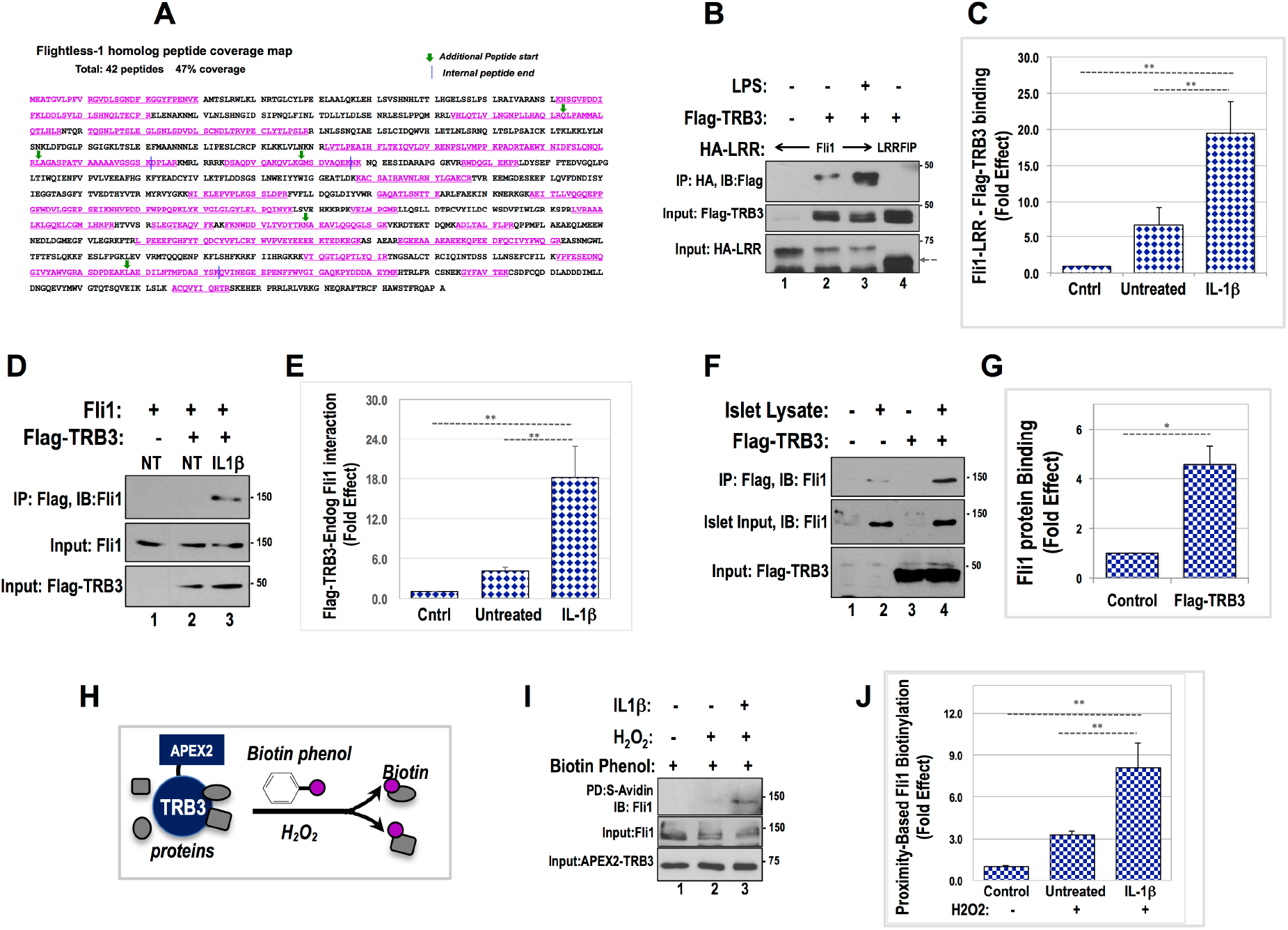
Cytokines induce TRB3 interaction with Fli1 (Flightless Holomog 1). **A)** Identification of Flightless-homolog 1 as a TRB3-interacting protein using mass spectrometry. The peptides identified are highlighted in pink font. Overlapping peptides are indicated with a green arrow for start site, and blue line denoting end of the internal peptide. Only peptides with peak intensity ratio of two-fold or higher on a Log2 scale were considered. **B)** Co-immunoprecipitation assay examines interaction between Fli1 and TRB3 proteins using HA-tagged LRR domain of Fli1 and Flag-TRB3, co-expressed in Min6 cells, without (lane 2) and following 30 min treatment with ILβ (lane 3). HA-Fli1-LRR in the absence of TRB3 served as control (lane 1) Note: TRB3 does not interact with HA-LRR-domain of LRRFIP2 (lane 4). **D)** Coimmunoprecipitation assay examines interaction between Flag-TRB3 and endogenous Fli1 from Min6 cells clustered into pseudoislets, without (lane 2) and upon 30 minute treatment with ILβ (lane 3). **F)** Protein interaction between purified Flag-TRB3 (lanes 3, 4) incubated with lysates from human islets (lanes 2, 4) showing potential for TRB3-Fli1 in human islets. **C, E**, **G**) Quantitative Analysis of data from Figures 2B, 2D and 2F depicted as fold induction of TRB3-Fli1 interaction over control. In each case, graphs represent average, fold-induction of TRB3-Fli interactions from three independent experiments each normalized to value of WT control (*\*\* p<0.005, * p<0.05)*. Schematic of proximity-induced biotinylation of APEX2 (ascorbic acid-peroxidase)-TRB3 interacting proteins. Peroxidase activity of APEX2-TRB3 fusion can cleave Biotin-phenol in the presence of H_2_O_2_ resulting in spontaneous biotinylation of proteins within 20 nm of TRB3. **I)** APEX2-TRB3 expressing Min6 cells were treated with Biotin-phenol (2 hours) followed by 1 minute of H_2_O_2_ treatment. IL1β stimulation was for 30 minutes prior to peroxide treatment (lane 3). Streptavidin-agarose was used to pull down biotinylated proteins and examined for presence of Fli1 using western blotting. Biotin-phenol without peroxide treatment was used as a control (lane 1). **J)** Fli1 recovered in western blots of streptavidin pulldowns was quantified and presented in graph format. Again, the data are represented as average fold-difference of Fli1 recovered by streptavidin beads in three independent experiments *(** p<0.005)*.

Next, we examined whether TRB3 could immunoprecipitate endogenous Fli1 from Min6 cells. Flag-TRB3 was expressed in Min6 cells, the cells treated with IL1β thirty minutes prior to harvesting. We conducted these experiments using TRB3 expressing Min6 cells that were clustered/ aggregated to form pseudo-islets. As shown in Figure 3D, under these conditions, we did not see appreciable interaction between TRB3 and endogenous Fli1 in untreated cells, but pretreatment with IL1β enabled Flag TRB3-Fli1 interaction. We also examined if TRB3 can interact with Fli1 in human islet extracts. We expressed and purified Flag-TRB3 from 293T cells, followed by purification using anti-flag antibody-linked agarose beads. The Flag-TRB3 immunoprecipitates were washed under high salt conditions to remove TRB3-bound endogenous Fli1 from 293T cells, before incubation with human islet protein extracts. Flag-antibody-crosslinked agarose beads were used as control. As seen in Figure 3F, TRB3 does interact with Fli-1 present in human islet extracts suggesting a potentially similar role for TRB3 in human islets.

Native, non-denaturing conditions such as those used to examine TRB3-interacting proteins in this study, can result in co-immunoprecipitation of direct protein interactors as well associated proteins in larger complexes not directly in contact with TRB3. We therefore assessed whether Fli1 protein interacts directly with TRB3 *in situ*. We used an assay that enabled proximity-dependent biotinylation of TRB3-interacting proteins. Briefly, this assay is premised on the fact that in the presence of H_2_O_2_, ascorbic acid peroxidase (APEX2) can activate biotin radicals using biotin phenol as substrate (see schematic Figure 3H). The short-lived biotin radicals thus generated are estimated to spontaneously attach to proteins within a range of 2-10 nanometers, which is about the width of an average protein.

Within such a short range, directly interacting proteins are most likely to be biotinylated. Based on this reasoning we examined whether APEX2-TRB3 fusion protein could biotinylate endogenous Fli1. The results of this assay are shown in Figure 3I. Cells expressing APEX2-TRB3 were treated with either biotin-phenol, or H_2_O_2_, or both, followed by streptavidin pulldown of biotinylated proteins. Notably, streptavidin pulldowns showed the presence of Fli1 protein only in the presence of both biotin-phenol and H_2_O_2_. TRB3-Fli1 interaction was enhanced in the presence of IL-1β. When taken together data from Figure 3 show that TRB3-Fli1 interacts with TRB3. Additionally, the use of a variety of approaches provides confidence in the authenticity of TRB3-Fli interaction.

### TRB3 disrupts Fli1-MyD88 interaction

In order for protein-protein interactions to be valid, they must satisfy appropriate functional parameters, and be consistent with the biological roles for each of the interacting proteins. Upon examination of the literature, we found that Fli1 has the ability to regulate TLR4 signaling, by binding and sequestering MyD88, a proximal and primary adaptor that recruits the downstream protein complexes required for IL1β-TLR4 signaling (35). It was reported that upon LPS stimulation, Fli1 dissociates from MyD88, freeing up MyD88 to participate in TLR4 signaling. Given that TLR4 and IL1β receptor both recruit MyD88 as the first adaptor (37), we examined whether MyD88-Fli1 interaction was impacted by TRB3 or by IL1β. To this end, we again used tagged proteins expressed in transfected Min6 cells, followed by co-IP assays. As seen in Figure 4A, under basal conditions, HA-Fli1 successfully immunoprecipitated Myc-MyD88. Published results have shown that LPS treatment can induce dissociation between Fli1 and MyD88 proteins (35). Similar to LPS, a 30-minute treatment with IL1β also markedly reduced HA-Fli1 pulldown of Myc-MyD88. Notably, co-expression of TRB3 strongly decreased Fli1-MyD88 interaction as well. These results suggest that TRB3 mimics IL1β-stimulated, post-receptor dissociation of Fli1 from MyD88. We have previously shown that TRB3 protein is nearly undetectable under basal conditions but is stabilized and detectable within 20-30 minutes following islet/Min6 cell stimulation with IL1β (5). This raises the intriguing possibility that IL1β-stimulated TRB3 induction could augment and prolong IL1β signaling by preventing premature re-association of MyD88 with the inhibitory Fli1 protein. If true, then the absence of TRB3 should provide a more permissive environment for Fli-MyD88 interaction even after IL1β stimulation.

**Figure 4:**
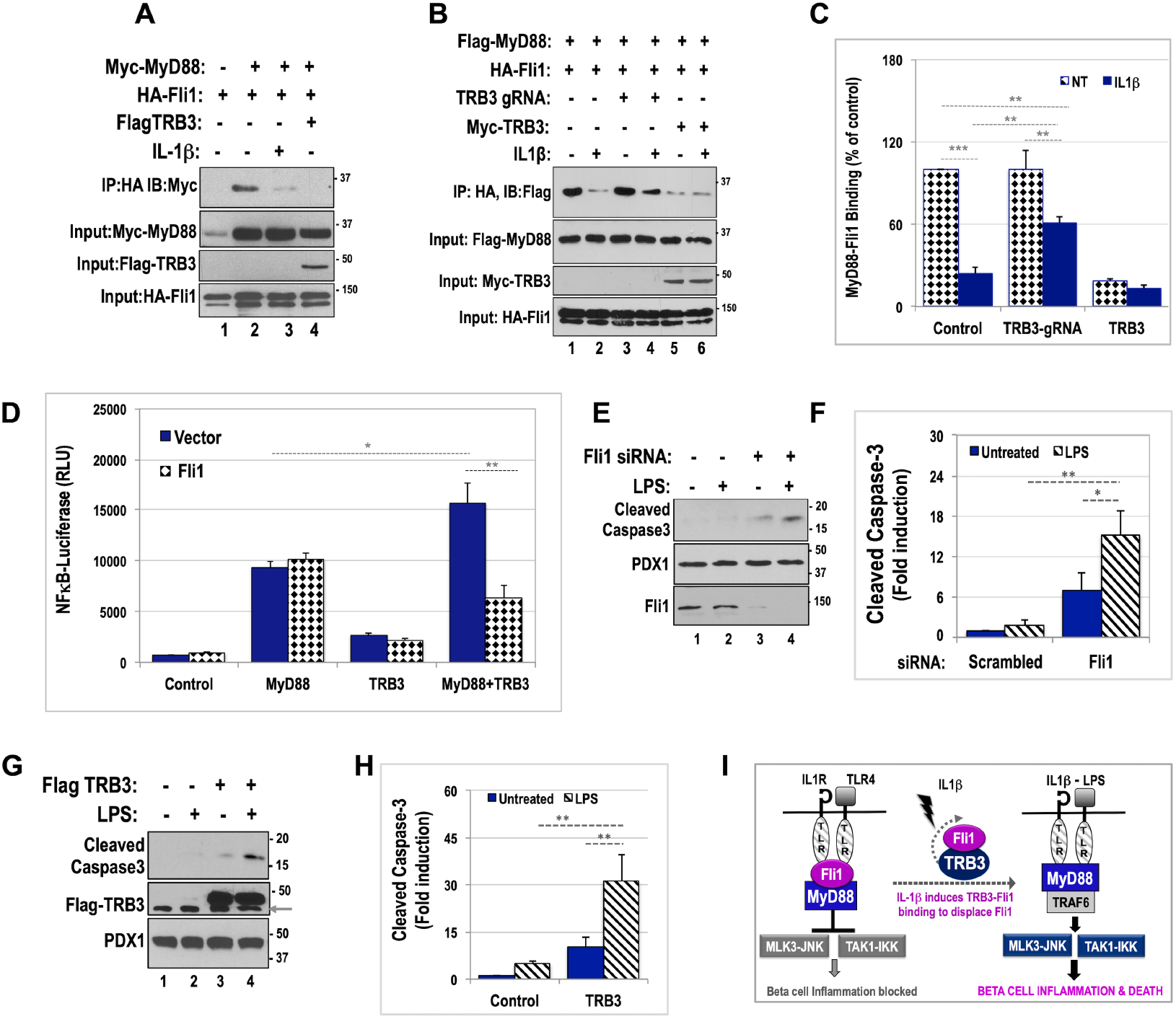
TRB3 disrupts Fli1-dependent anti-inflammatory sequestration of Myd88. **A)** Protein interaction between Fli1 and MyD88 proteins was assessed using lysates from Min6 cells transfected with HA-Fli1 (1–4) and Myc-MyD88 (lanes 2-4) in the presence of control vector (lane 1), upon 30 min IL1β treatment (lane 3), or expression of HA-TRB3 (lane 4). Inputs for each protein are shown **B)** Impact of TRB3 knockdown on Fli1 and MyD88 protein interaction was assessed using identical conditions as in Figure 4A, in the presence of control gRNA, (lanes 1-2), TRB3 gRNA (lanes 3-4) and Myc-TRB3 (lanes 5-6), and in the presence of vehicle (lanes 1, 3 and 5) or upon 30 minute pretreatment with IL1β. Of note, TRB3 knockdown results in partial retention of Fli1-Myd88 interaction even in the presence of IL1β (compare lanes 2 and 4). **C)** Fli1-Myd88 interactions from Fig 4B were quantified, and amount of MyD88 recovered was expressed as a fraction of MyD88 input. Stippled and solid bars denote Fli1-MyD88 binding in the absence or presence of IL1β respectively. Two-way ANOVA was performed to compare WT and TRB-gRNA (\*\**p<0.05*, \**\*\*p<0.005)* **D)** Luciferase reporter assay from Min6 insulinoma cells transiently transfected with NFkB-Luciferase (firefly) reporter in the presence of MyD88, TRB3 or both, in the absence (solid bars) or presence (stippled bars) of Fli1. Data presented are an average of triplicate values +/−SD from a representative experiment (n=5). Two-way ANOVA of TRB3 expression on MyD88 effect (\**p=0.05, **p=0.005)* **E)** LPS treatment (lanes 2 and 4) enhances Fli1-siRNA (lanes 3-4) knockdown-mediated cell death as judged by increasing expression of cleaved caspase 3 (compare lanes 3 and 4). LPS treatment alone (lane 2) does not induce cell death in the presence of scrambled siRNA (lanes 1-2). **G)** Adenovirus-mediated TRB3 overexpression (72 hrs-lanes 3-4) induces low-level cell death (lane 3), which is enhanced by 48 hr LPS treatment. LPS treatment alone (lane 2) does not induce cell death/caspase3-cleavage in the presence of control virus (lanes 1-2). **F and H**) Fold induction of cleaved-caspase3 was calculated using densitometric analysis of data from three independent experiments, and analyzed using two-way ANOVA *(*p=0.05, **p=0.005)*. **I)** Schematic model summarizing data presented and depicting mechanism by which TRB3 engages Fli1 to impact IL1/TLR4 signaling.

We examined this question by using Fli1-MyD88 co-IP assays similar to those in Figure 4A, this time using an excess of TRB3 or knockdown of TRB3 expression. As seen in Figure 4B, HA-Fli1 (co-transfected with control gRNA), showed strong binding to MyD88 under basal unstimulated conditions (quantified and arbitrarily assigned as 100% binding in Figure 4C), which was markedly reduced to roughly 20% upon IL1β stimulation for 30 minutes. We used TRB3 gRNA for knockdown of TRB3 in the same assay and found that as expected, knockdown of TRB3 did not significantly impact strength of basal, unstimulated Fli-MyD88 interaction. However, upon IL1β stimulation, rather than an expected 80% drop in Fli1-MyD88 interaction, knockdown of TRB3 resulted in a only 40-45% drop in Fli1-MyD88 interaction (Fli1-MyD88 interaction quantified in Fig 4C, corrected for expression levels of proteins). Unsurprisingly, co-expression of TRB3 severely abrogated both basal and IL1β mediated Fli1-MyD88 interaction. Taken together these data show that in the absence of TRB3, Fli1 retains partial ability to sequester MyD88 and thereby attenuate IL1β-dependent signaling, while presence of TRB3 could augment IL1β signaling by overcoming Fli1-dependent brake on MyD88 recruitment.

### TRB3 enhances MyD88-dependent activation of NFkB reporter

In Figure 2 we showed that TRB3 gRNA is sufficient to blunt IL1β-dependent activation of NFkB-luciferase. Considering that the robustness of IL1R/TLR driven signaling can be modulated by availability of MyD88, and that the presence of TRB3 creates permissive conditions for MyD88 recruitment to IL1/TLR receptors, we asked whether overexpression of MyD88 was sufficient to activate NFkB-dependent transactivation and whether TRB3 could impact such an effect of Myd88. As shown in Figure 4D, overexpression of MyD88 strongly activated NFkB-luciferase activity (>10 fold), even in the absence of upstream receptor stimulation. While TRB3 alone did not significantly alter NFkB-luciferase activity, TRB3 significantly potentiated MyD88-dependent NFkB-luciferase activity increasing it from 10-fold activation to a 15 to17-fold effect (compare dark blue bars from L-R in Figure 4D). Interestingly, Fli1 co-expression (represented by hatched bars across the figure) did not impact MyD88-dependent, NFkB-luciferase activity. These data are consistent with the fact that Fli1 mediated sequestration of MyD88 is supported by additional proteins or factors and the stoichiometry of endogenous interactions is most likely overcome by excess of expressed proteins in this artificial but informative assay. It is particularly noteworthy that while Fli1 does not impact MyD88 activity under these conditions, Fli1 co-expression is sufficient to abrogate TRB3-mediated potentiation of MyD88-activated NFkB-luciferase activity. We interpret these data to suggest that the ability of TRB3 to potentiate IL1β/TLR/MyD88 signaling could depend directly on its ability to bind Fli1, rather than a complex of Fli1-associated factors.

### Fli1 knockdown, and TRB3 overexpression augment LPS-induced cell death

We have thus far seen that TRB3 influences inflammation at least in part by binding and disabling Fli1 which serves as a brake on IL1/TLR signaling in the absence of an appropriate stimulus. We examined whether Fli1 loss of function was sufficient for abnormal activation of inflammatory cell death. To overcome the high expression and long half-life of Fli1, we performed dual transfection of Fli1 siRNA in which cells are transfected with Fli1 or control siRNA for one passage (3-4 days) prior to, as well as during the actual experiment. Such a regimen was required for the significant knockdown of Fli1 as seen in Figure 4E. 72 hours after the second siRNA transfection, cells were treated with TLR4-specific LPS, harvested after 48 hours and examined for expression of cleaved caspase3, an indicator of end-stage cellular apoptosis. Interestingly, LPS-treatment of Min6 cells even for 48 hours, showed negligible expression of cleaved caspase3 (Figure 4E) perhaps because LPS activates IKK, but does not activate the JNK pathway in beta cells. Chronic knockdown of Fli1 resulted in a low-level induction of cleaved caspase 3, which was significantly augmented by the presence of LPS.

We have shown in Figure 2K that islets from TRB3KO mice, can confer protection from pro-inflammatory effects of cytokines. We examined whether gain of TRB3 function could accelerate inflammatory cell death. Min6 cells were infected with adenovirus encoding Flag-tagged TRB3, treated with LPS for 48 hours and lysates probed for presence of cleaved caspase3. TRB3 expression alone is sufficient to induce cell stress, it was unclear if it can induce cell death in the absence of an pro-inflammatory milieu. Again, we used LPS treatment to examine whether presence of TRB3 can augment inflammation and cell death. Expression of TRB3 resulted in low-level increase in caspase3 cleavage, but addition of LPS strongly augmented cleaved caspase3 expression, suggesting that TRB3 can potentiate inflammation and accelerate beta cell apoptosis. While data presented in Figures 4E and 4G are interesting, particularly noteworthy is the striking similarity between Fli1 knockdown and TRB3 overexpression to induce low level cell death and the ability of LPS to augment apoptosis in both cases. These data are consistent with the ability of TRB3 to engage and inhibit Fli1 action. Finally, in Figure 4I, we present a schematic depiction for the potential mechanism by which TRB3 boosts inflammatory signaling. The model also offers an explanation for resistance of TRB3KO islets to cytokine-induced beta cell death. We conclude that TRB3 and Fli1 have reciprocal roles in beta cell inflammation and in cytokine-induced cell death.

## Discussion

The role of TRB3 in ER stress has been extensively examined. However, its role in inflammation is less well documented. In this study we have examined the mechanism by which TRB3 supports inflammatory stress in beta cells. We have previously shown that IL1β and conditions mimicking insulitis induce TRB3 protein expression in both mouse and human islets (5,10). In response to IL1β, TRB3 binds and stabilizes MLK3 an upstream MAP3K to maintain activation of the JNK module and drive a full and strong inflammatory response, (5) such that islets from TRB3 null mice show a roughly 50% decline in overall activation of MLK3-JNK pathway (10). In addition to MLK3-JNK, a full-blown inflammatory response in beta cells also requires activation of a second arm of signaling driven by TAK1-IKK-NFkB pathway (15). In this study we report that similar to MLK3-JNK, phosphorylation and activation of TAK1-IKK pathway is also attenuated in TRB3KO islets, resulting in partially suppressed NFkB-dependent gene expression and an overall dampening of the proinflammatory response. At the mechanistic level we find that unlike the MLK3-JNK module, TRB3 does not directly engage TAK1 or IKK, suggesting a more upstream role for TRB3. Here we have reported a novel interaction between TRB3 and Fli1, an immuno-modulatory, actin-binding protein that controls the strength of post-receptor IL1/TLR4 signaling. Specifically, Fli1 sequesters MyD88, a crucial and proximal adaptor that is recruited to the toll-like-receptor (TLR) domain at the cytoplasmic end of both IL1β and TLR4 receptors (35). Upon receptor recruitment, MyD88-forms a complex with IRAKs1 and 4 (IL receptor associated kinase), triggering cross-phosphorylation of IRAKS, ultimately leading to TRAF6-mediated ubiquitination and activation of specific MAP3Ks including TAK1 (37). Consistent with the ability of TLR4 and IL1β to signal via shared mechanisms, we find that TRB3 modulates both IL1β and TLR4-dependent IKK activation.

IL1β is one of the most potent cytokines for inducing inflammation in the beta cell (38). By contrast, TLR4 is a pattern recognition receptor that recognizes pathogen associated molecular patterns (PAMPs) and presents a first line of defense against pathogens by binding lipo-polysaccharides (LPS) from bacterial remnants (39). TLR4 activation is required for saturated fatty acid-mediated inflammation and dysfunction (22). Saturated fatty acids like palmitic acid are crucial agents for the slow and prolonged inflammatory process of lipo-glucotoxicity that is central to beta cell inflammation in T2D (21,40). Interestingly, TLR4 signaling in beta cells showed little activation of JNK (not shown), and a slightly delayed TAK1-IKK-NFkB activation profile, perhaps explaining the slower developing profile of lipotoxicity. Indeed, TLR4KO and MyD88KO mice are resistant to saturated fatty acid-induced islet dysfunction and inflammation (22). Despite the demonstrated expression and of TLR4 on beta cells(41–43), there remains some ambiguity regarding the ability of LPS to directly stimulate beta cells. A second school of thought subscribes to the belief that TLR4-stimulated signaling of islet beta cells is indirect, and a result of secondary release of IL1**β** from LPS-stimulated, islet-resident macrophages. These conclusions were based on beta cells from disaggregated islets. The discrepancy between the two schools of thought could be reconciled by the potential role of macrophage dependent release of CD1 and MD2 co-receptors (44,45) required for LPS-dependent TLR4 activation and deserves further examination.

TAK1-IKK-NFkB pathway is central to cellular inflammatory responses. A complete inhibition of NFkB would be expected to inhibit all target genes. However, our results show that loss of TRB3 attenuates but does not abolish IKK or JNK. Interestingly, such a decrease in peak activity and temporal activation of IKK-NFkB does not uniformly impact downstream gene expression targets (1,46). Our findings are consistent with the discovery that modulation of amplitude and/or duration of kinase activation itself encodes key information for driving distinct gene expression programs. Complexity of promoters, coupled with cooperative gene activation by transcription factors that are networked and integrated with upstream signals, likely renders each gene uniquely responsive to kinetics of IKK activation. In the beta cell, key NFkB gene targets iNOS, proinflammatory cytokines and chemotaxis promoting protein MCP1 (macrophage chemoattractant protein)/CCL2 activate a feed forward inflammatory loop between islet resident macrophages and beta cells (47,48). TRB3KO islets displayed a sharp decrease in MCP1/CCL2 gene expression, and a comparatively modest decrease in iNOS gene activation. Surprisingly, we observed a small (less than 20%) decrease in MCP1 protein expression (not shown) but near complete inhibition of iNOS protein expression in response to IL1β and TLR4 stimulation. These data highlight, and future studies will examine the potential role for TRB3 in iNOS protein expression. Our data show a crucial role for TRB3 in post-transcriptional regulation of cytokine-activated iNOS and MCP1 expression and serve as a caution against over-reliance on mRNA data to explain biological responses.

Our data demonstrate that TRB3 influences iNOS expression at the transcriptional as well as a post-transcriptional level. NFkB-driven iNOS expression is crucial for cytokine and saturated fatty acid-induced beta cell damage (29). Depending on the cell type, iNOS dependent nitric-oxide (NO) production is associated with effects ranging from increased survival to favoring cell death. In cells where it has a deleterious role, NO generates powerful peroxynitrites, which augment oxidative stress and accelerate inflammation-induced cell damage. (49,50). A myriad of pleiotropic activities of NO has precluded a real consensus on the mechanism by which NO regulates cytokine-activated cell death (51,52). Liu et al showed that compared with control islets, primary islets from INOS−/− mice display 50-80% decrease in cytokine-induced necrosis (53), while Holohan et al showed that overexpression of iNOS alone is sufficient to induce beta cell apoptosis (54). We conclude that a strong inhibition of iNOS expression in TRB3KO islets may at least partially explain their resistance to cytokine-induced apoptosis.

TRB3 has been demonstrated to modulate kinase action by direct interaction with several kinases including mTORC1, AKT, MLK3 and MAP2Ks (3,5,9,55). Here we identify TRB3-Fli1 interaction as a novel mechanism by which TRB3 augments beta cell inflammation and death. Fli1 can exert a synergistic or an antagonistic inflammatory role in a context dependent manner (13) (56). Nevertheless, the rapidly expanding proinflammatory role of Fli1, lends credence to the ability of TRB3 to augment cytokine-dependent beta cell stress via Fli1. Results reported in this study suggest that TRB3 and Fli1 function in a reciprocal fashion. Fli1 and TRB3 behave like scaffold proteins to introduce plasticity into the signaling process. Neither the gain of TRB3 function, not the loss of Fli1 by themselves are sufficient to prompt rapid activation of cell death. In each case, LPS which itself has very pro-apoptotic effects, was required to induce death suggesting that both Fli1 and TRB3 modulate rather than completely alter the dynamics of inflammation. Scaffold-to-scaffold interactions are increasingly being recognized as highly dynamic networks that control multiprotein complex formation. These transient interactions alter the temporal and kinetic flow of cellular information, to elicit distinct biological and metabolic outcomes (57–59). Based on its ability to bind Fli1 and modulate the dynamics of receptor recruitment of MyD88, our results recast TRB3 as a protein that can modify flow of information to produce distinct biological outcomes.

## Experimental Procedures

### TRB3 Knockout mice

The mutant mouse strain was obtained from the European Mouse Mutant Archive (EMMA ID EM:02346), Helmholtz Zentrum Muenchen, German as described (10). Mice were housed in a 12-h light/12-h dark cycle at controlled temperature (25 °C ± 1 °C). Genotyping was performed by PCR of genomic DNA according to above referenced protocols. In experimental procedures, we compared homozygous TRB3 knockout mice with wildtype littermates (control mice). TLR4KO mice were provided by Dr. Richard Gallo (Dept. of Dermatology, UCSD). All experimental procedures were performed according to University of California San Diego IACUC policies.

### Reagents

Antibodies used for western blotting included anti-Myc, anti-Flag, anti-HA, anti-pJNK, anti-JNK anti-pIKK, anti-TAK1, anti-pTAK1, Cleaved Caspase-3, anti-β-tubulin from Cell Signaling, Beverly, MA. Anti-Fli1, and anti-iNOS were purchased from Santacruz Biotechnology, Dallas TX, and anti-TRB3 antibodies were a gift from Dr. Marc Montminy, Salk Institute, CA. For immunofluorescence, mouse anti-BAX clone 6A7 (BD Bioscience, San Jose, CA), sheep anti-insulin (Binding Site, CA), and In Situ Cell Death Detection Kit (Fluorescein) from Roche (Indianapolis, IN) were purchased from commercial sources. Agarose-conjugated antibodies used for immunoprecipitations included Anti-HA agarose (Thermo-Fisher Scientific, Waltham, MA), and Anti-FlagM2 agarose (Sigma-Aldrich, St. Louis, MO). Streptavidin-Agarose was purchased from Pierce. Immunofluorescence and western blotting employed fluorescent or HRP-conjugated secondary antibodies (Jackson ImmunoResearch, West Grove, PA), the latter detected using Supersignal West-Pico Plus chemiluminescence reagent (Thermo-Fisher, Waltham, MA). Other reagents include IL-1β, (Peprotech, Rocky Hill, NJ) and ultrapure TLR4-specific LPS (Invivogen, San Diego CA). Flag-TRB3 adenovirus was generated as described 3). Smartpool TARGETplus siRNA was used to knockdown Fli1, and non-targeting siRNA was used as control (Dharmacon-Horizon Discovery Sciences). TPCA-1 (Bio-Techne-Tocris, Minneapolis, MN) was used to inhibit NFkB pathway, and MLK3 was inhibited using CEP11004 (gift from Cephalon Inc.)

### Plasmids and Constructs

HA-TRB3, FLAG-β-tubulin, constructs have been described elsewhere (5). Plasmids encoding HA-tagged LRR (leucine-rich-repeat) domains of LRRFIP2 (Addgene plasmid # 21152), and Fli1 (Addgene plasmid # 21151), were a gift from Dr. Junying Yuan (Harvard University, Boston MA), and Flag-MyD88 (Addgene plasmid # 13093) was a gift from Dr. Ruslan Medzhitov (HHMI, Yale School of Medicine). pcDNA3-HA-Fli1 was generated by PCR-based amplification of full-length, mouse Fli1, using pCMV-Sport6 mFli1 (Horizon Discovery, Denver CO) as template, and cloned into pcDNA3.1 vector using HindIII and Xho1 sites, in frame with a C-terminal HA tag. Myc-APEX2-TRB3 was generated by PCR amplification of APEX2 using pcDNA3-APEX2-NES as template (Addgene plasmid #49386, kind gift of Alice Ting, Stanford University, Palo Alto, CA), and subcloned into BamH1 and Xho1 sites of pcDNA3 Myc plasmid, followed by subcloning of TRB3 in frame with APEX2, using EcoRI and XbaI sites). NFkB-luciferase was purchased from Clontech. Nickase-Ninja TRB3 gRNA plasmids were purchased from ATUM Bioscience (Newark, CA). Each plasmid expressed CMV promoter driving a Cas9N nickase mutant that causes single strand breaks, and co-expresses two gRNAs using dual U6 promoters. gRNA directed against GFP was included as control (60). Sequence of dual gRNAs targeting either Exon 2 or Exon 3 of the mouse TRB3 gene, are included in Supporting Information section.

### Primary islet isolation and Cell Culture

Human islets were obtained from the Integrated Islet Distribution Program. Islets were hand-picked and cultured overnight in CRML supplemented with 10% FBS. Prior to experiments, islets were cultured in RPMI-1640, with 11 mM glucose and supplemented with 10% FBS. Mouse islets from WT and TRB3KO mice were isolated as described, using Liberase (0.2 mg/ml; Roche Applied Science) to distend the pancreas, and digested at 37 °C for 15 min. Preparations were washed with Hanks’ buffered salt solution, and dissociated islets purified using Histopaque (Sigma Aldrich) gradients and cultured in RPMI 1640 medium supplemented with 10% FBS. Min6 cells (passages 15-18 *only*) were grown in DMEM containing 25mM glucose supplemented with 4% heat inactivated FBS and 50μM β-mercaptoethanol.

### Transient Transfections, Nucleofection and Luciferase reporter assays

Transfections for reporter assays and co-IP experiments were carried out using Lipofectamine 2000 (Invitrogen) as per manufacturer’s instructions and 40-48 hours post-transfection, cells were treated with 10ng/ml IL-1β, or 10ng/ml TLR4-specific LPS as described. For gRNA experiments, 1×10^6^ Min6 cells were electroporated using Nucleofector Kit V (Amaxa GmbH, Cologna, Germany) with 0.5-1μg of plasmid DNA encoding TRB3-targeted gRNA (described above), and 0.4μg of pcDNA3-RFP as nucleofection control and used for analyses 40-48 hours post-nucleofection. Luciferase assays were performed using CSH Protocols as described. For Guassia luciferase, 10-30 ul of secreted medium was used with coelentrazine as substrate. Luciferase activity was measured using a tube luminometer (Berthold Technologies). For efficient knockdown of Fli1, a long-lived protein, dual siRNA transfections were performed using Dharmafect-3 reagent at a final concentration of 50 nm. Cells intended for Fli1 knockdown experiments were pre-transfected with respective siRNAs, split after 72-96 hours, and re-transfected with the same siRNAs before proceeding to conduct the experiments after a few days as described.

### RNA Isolation and Q-PCR

Islets isolated from WT and TRB3KO mice were stimulated with IL-1β for 2 and 4 hours. Total RNA was isolated using RNAqueous isolation kit (Thermo-Fisher, Waltham MA) and mRNA reverse transcribed using PrimeScript RT Reagent Kit (Takara Bio USA, Mountain View CA). Quantitative PCR reactions were carried out using PowerUp SYBR-Green master mix using StepOne Real-Time PCR systems (Applied Biosystems). GAPDH mRNA levels were used for normalization. Relative mRNA expression was determined by ΔΔCT method. Primers used for qPCR reactions are listed will be provided on request.

### Proximity-Dependent Biotinylation of TRB3-interacting proteins

10 cm dishes plated with 8-10 million Min6 cells were transfected with 8 ug of plasmid expressing APEX2-TRB3 fusion protein. 40-48 hours post-transfection, the cells were treated with 2 mM Biotin-tyramide (APEX-Bio Inc) for 2 hours. Biotinylation was activated for exactly 1-minute using a final concentration of 0.5mM H_2_O_2_, and terminated using quenching solution as described (61). Biotinylated proteins in cellular lysates were pulled down using streptavidin-agarose, resolved on SDS-PAGE and probed for interacting proteins using western blotting.

### Splenocyte and Islet Co-Culture (SICC)

<u>Mouse SICC</u> was performed as previously described ((10,18)). Briefly, wild-type spleens were crushed and passed through a 70μm mesh. Red blood cells were lysed in 0.15M NH4Cl and 1.5 × 10^6^ splenocytes per well were plated in 24 well dishes in RPMI supplemented with 10% heat-inactivated FBS. Splenocytes were stimulated with plate bound anti-CD3 and exogenous anti-CD28 antibodies (10μg/ml and 1μg/ml respectively; BD Biosc, San Diego, CA) for 2 days. Isolated islets from WT or TRB3KO mice were rested overnight, and co-cultured using transwell filters in the presence of unstimulated or stimulated splenocytes. <u>Human SICC</u> was done using similar protocol slightly modified protocol as described (18). For drug treatments, CEP11004 was provided by Cephalon Inc, TPCA1 and BAY-11-1085 were purchased from Tocris Inc.

### Immunofluorescence and TUNEL assays, microscopy and image acquisition

Islets were fixed in Zinc buffered formalin (Anatech, Battle Creek, MI) for 1 hour and processed for routine cryo-microtomy. TUNEL assay was carried out according to manufacturers instructions (Roche Diagnostics, IN). Primary antibodies were visualized with species-specific secondary antibodies conjugated to fluorescent probes. Fluorescent images were acquired on a Zeiss 710 confocal microscope. All images were assembled in Photoshop CS6, (Adobe Systems Inc., San Jose, CA) and ImageJ (NIH). DAPI (1μg/ml) was used as a nuclear counterstain.

### Protein-protein Interactions & Co-immunoprecipitations

For identification of TRB3 interaction Proteins, freshly split Min6 cells (passage 16) were infected with Flag-TRB3 adenovirus or HA-tagged TRB3 lentivirus and approx. 100×10^6^ cells in multiple batches of 10-12×10^6^ cells were allowed to form pseudo-islet clusters with gentle shaking in the cell-culture incubator and medium replaced after 16-20 hours. Cells/pseudo-islets were pooled and harvested at 36 hours, using standard co-IP conditions using NEHN buffer containing 150 mM NaCl, 0.5mM EDTA, 20mM HEPES pH 7.8, and 0.5% NP-40 (Pierce), with a full complement of protease and phosphatase inhibitors. Lysates were subjected to immunoprecipitation using precleared agarose-bound Flag or HA tag-specific antibodies for 4 hours and eluates washed 5 times with gentle shaking, for 3-5 minutes, resolved using SDS-PAGE, stained using colloidal Coomassie stain, destained, specific bands excised and subjected to in-gel trypsinization followed by mass-spectrometric analysis. GFP adenovirus or RFP lentivirus were used as control. The experiment was repeated using Min6 cells. Pseudo-islets were used for co-IP experiments of IL1β induced TRB3 binding to endogenous Fli1. Min6 monolayers were used for interaction of expressed proteins. For TRB3 interaction with Fli1 from human islets, Flag-tagged TRB3 was expressed in cultured cells, purified by immunoprecipitation using NEHN, the beads washed with NEHN-twice with 300 mM NaCl αconcentration and twice times with NEHN-150 mM NaCl.

### Sample preparation for mass spectrometry

Sample immunoprecipitates were extracted in guanidine solution, methanol precipitated, and pellet suspended 8M Urea-Tris buffer followed by 10 mM TCEP mM and 40 mM chloro-acetamide solution, the urea concentration reduced to 2M and used for tryptic digestion. Digested peptides were acidified (0.5% TCA), desalted and concentration measured using BCA. 1 ug proteins were injected for each label-free quantification run.

### Mass-Spectrometry and identification of proteins

Trypsin-digested peptides were analyzed by ultra-high pressure liquid chromatography (UPLC) coupled with tandem mass spectroscopy (LC-MS/MS) using nano-spray ionization. The nanospray ionization experiments were performed using a Orbitrap fusion Lumos hybrid mass spectrometer (Thermo) interfaced with nano-scale reversed-phase UPLC (Thermo Dionex UltiMate™ 3000 RSLC nano System) using a 25 cm, 75-micron ID glass capillary packed with 1.7-μm C18 (130) BEH™ beads (Waters corporation). Peptides were eluted from the C18 column into the mass spectrometer using a linear gradient (5–80%) of ACN (Acetonitrile) at a flow rate of 375 μl/min for 1h. The buffers used to create the ACN gradient were: Buffer A (98% H2O, 2% ACN, 0.1% formic acid) and Buffer B (100% ACN, 0.1% formic acid). Mass spectrometer parameters are as follows; an MS1 survey scan using the orbitrap detector (mass range (m/z): 400–1500 (using quadrupole isolation), 120,000 resolution setting, spray voltage of 2200V, ion transfer tube temperature of 275 °C, AGC target of 400,000, and maximum injection time of 50ms was followed by data dependent scans (top speed for most intense ions, with charge state set to only include+2–5 ions, and 5s exclusion time, while selecting ions with minimal intensities of 50,000 at which the collision event was carried out in the high-energy collision cell (HCD Collision Energy of 30%), and the fragment masses were analyzed in the ion trap mass analyzer (with ion trap scan rate of turbo, first mass m/z was 100, AGC Target 5000 and maximum injection time of 35ms). Mass-spec parameters were as described. Protein identification and label free quantification was carried out using Peaks Studio 8.5 (Bioinformatics solutions Inc.)

### Statistics

Where applicable, area under the curve (AUC) values were calculated using GraphPad Prism software. Differences between means were examined using analysis of variance (ANOVA) followed by a Bonferroni *post hoc* comparison were also performed using GraphPad Prism software. Unless specified, data presented in bar graphs show average fold effects from three independent experiments. Within each experiment values were normalized to WT control, assigned a value of 1.

## Supporting Information

**Figure S1A:**
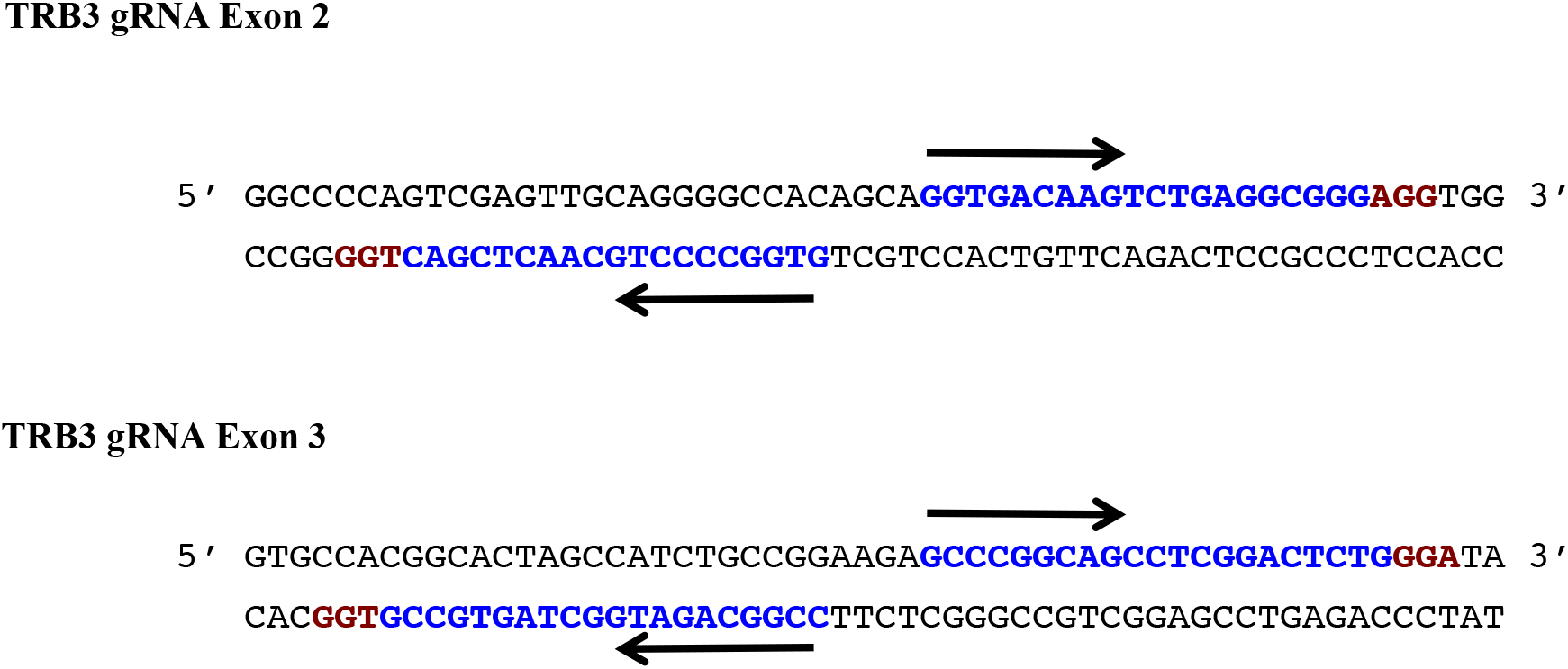
Sequence and configuration of TRB3 gRNA for Cas9 D10A Nickase based genome editing: Design of two independent gRNAs targeting Exon 2 (A) and Exon 3 (B) of the TRB3 gene. The PAM sequences are highlighted in red and two gRNA are highlighted in blue. Arrows indicate the orientation of the oligomer/gRNA is indicated by black arrows.

**Figure S2A:**
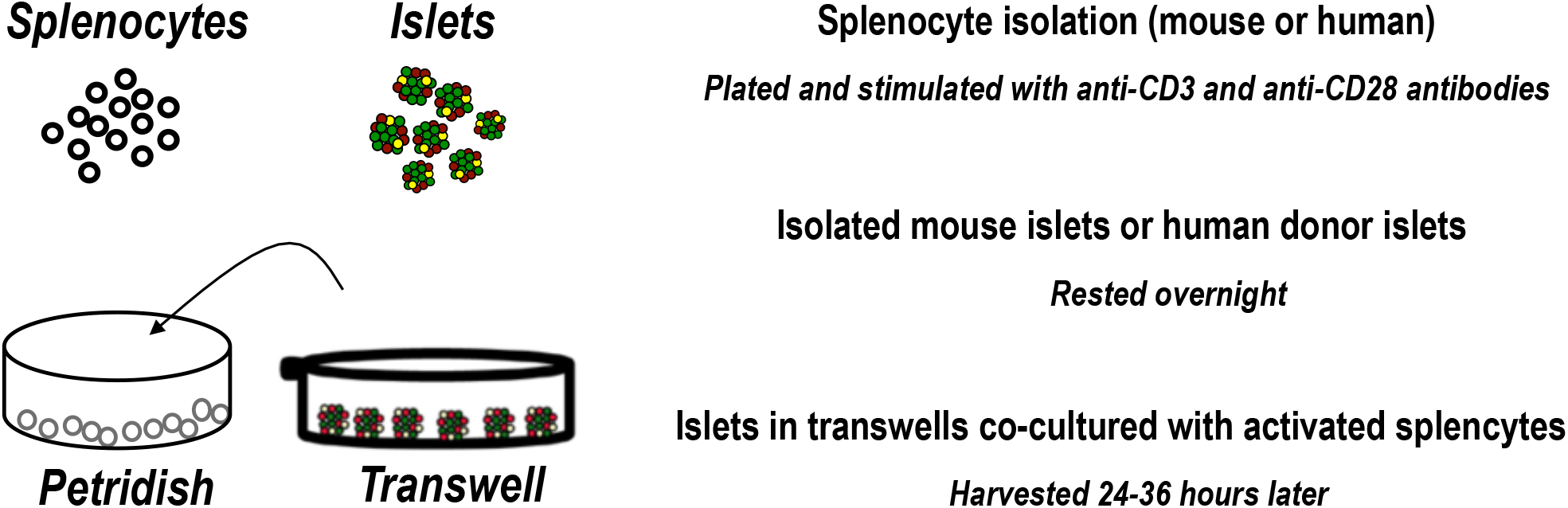
Schematic depicting Splenocyte-Islet co culture: Primary splenocytes from WT mice (or cadaveric human donor) were isolated on day 1 and plated as indicated to activate an immune response. Human donor islets or islets from WT and TRB3KO mice, were isolated the next day and allowed to rest and each set of islets was cocultured with activated or unstimulated splenocytes on day 3. For human splenocytes, cadaveric human spleen was used.

**Table S1:** Secretory profile of stimulated splenocytes: Medium secreted by unstimulated and stimulated splenocytes from human splenocytes (**A**) or mouse splenocytes (**B**) was profiled for the indicated cytokines by multiplex ELISA.

|  | Unstim | Stim 48 hrs | Unstim | Stim 48 hrs |
| --- | --- | --- | --- | --- |
| **TNF-α** | **3** | **46.6** | **167** | **4385** |
| **IL-4** | **3** | **137** | **12** | **1181** |
| **IL-5** | **3.5** | **68.3** | **7** | **345** |
| **IL-6** | **7.3** | **404** | **1825** | **4108** |
| **IL-10** | **6.99** | **333** | **97** | **9030** |
| **RANTES** | **27** | **191** | **150** | **8860** |
| **GM-CSF** | **3.2** | **370** | **219** | **5366** |
| **IFN-γ** | **4.75** | **10200** | **725** | **8690** |
| **IL-1β** | **1.2** | **13.6** | **51** | **595** |

| sample ID | Wells | GM-CSF<br>pg/mL | IFN-g<br>pg/mL | IL-10<br>pg/mL | IL-1β<br>pg/mL | IL-6<br>pg/mL | MCP-1<br>pg/mL | MIP-1a<br>pg/mL | Rantes<br>pg/mL | TNF-a<br>pg/mL |
| --- | --- | --- | --- | --- | --- | --- | --- | --- | --- | --- |
| WT Stim 24hr | G8 H8 | 43.1 | 400 | 32.9 | <3.2 | 179 | 28.3 | 485 | 35.3 | 20.4 |
| WT Stim 48hr | A9 B9 | 1570 | 12000 | 685 | 3.87 | 973 | 149 | 9050 | 115 | 133 |
| WT Stim 72hr | C9 D9 | 2610 | 12100 | 1760 | 4.55 | 1240 | 400 | 9200 | 142 | 145 |
| WT unstim 24hr | C10 D10 | <3.2 | <1.25 | 7.94 | <3.2 | 36.8 | <3.2 | 71.2 | 23.7 | 1.91 |
| WT unstim 48hr | E10 F10 | 9.12 | <1.25 | 12.5 | <3.2 | 57.2 | <3.2 | 75.9 | 30.4 | 3.86 |
| WT unstim 72hr | G10 H10 | <3.2 | <1.25 | 5.65 | <3.2 | 50.1 | <3.2 | 58.2 | 27.2 | 1.19 |

